# Solving the gene classification problem with a novel Gene Space approach

**DOI:** 10.1101/2025.10.17.683012

**Authors:** Konstantin S. Zaytsev, Natalya S. Bogatyreva, Alexey N. Fedorov

## Abstract

Genomic organization and its comparative analysis throughout all major kingdoms of life are extensively studied across multiple scales, ranging from individual gene-level analyses to system-wide investigations. This work introduces a novel framework for characterizing genetic architecture through a new integral genomic parameter. We propose the concept of a multidimensional Gene Space to enable holistic quantification of genome organization principles. Gene Space — a multidimensional space based on the frequencies of nucleotide tokens, such as individual nucleotides, codons, or codon pairs. We demonstrate that in this space, genes from each of the studied microorganism species occupy a limited region, and individual genes from different species can be effectively separated. Consequently, a specific Genome Subspace can be defined for each species, which constrains the organism’s evolutionary pathways, thereby determining the constraints on gene optimization for these species. Further in-depth analysis is required to test if it is true for other organisms as well. The Gene Space framework offers a novel and powerful approach for genome analysis at the most basic levels, with promising applications in comparative genomics, evolutionary biology, and gene optimization.

## INTRODUCTION

The genetic code is a universal apparatus essential to all living organisms. Each codon determines which amino acid will be incorporated into the synthesized protein. With 61 codons encoding 20 distinct amino acids, the genetic code is redundant: the same amino acid can be encoded by one to six synonymous codons. Consequently, numerous nucleotide sequences can encode identical amino acid sequences.

Despite this diversity, organisms have preferences for specific nucleotide combinations. For example, genes from different organisms encoding homologous proteins have significant differences in codon usage bias: in every organism, certain codons are used more frequently than others, forming species-specific codon usage profiles [1,2]. A key driver for these differences is the GC content [3,4], which describes the difference in usage of A+T and G+C nucleotides [5].

During protein synthesis, each codon is recognized by a specific tRNA. Thus, codon frequencies in the genome correlate with the abundance of corresponding tRNAs [6,7]. Codon bias directly impacts translation efficiency: overuse of a codon can deplete its cognate tRNA, reducing overall protein synthesis efficiency and affecting organismal viability [8].

In recombinant gene expression, the primary goal is maximizing target protein yield. However, disrupting cellular process balance may induce cell stress [9]. Therefore, a common gene optimization strategy involves adjusting codon frequencies to match host organism preferences. Crucially, not only individual codon frequencies but also their combinations (correlations between adjacent codons) influence protein synthesis efficiency [10].

Optimization typically is based on genome-wide averaged codon bias. Yet codon bias varies significantly not only among genes but also within different regions of the same gene [2,11]. Notably, genes with varying expression levels exhibit distinct codon biases [2,12]. Our previous research further demonstrates that codon frequency optimization characterizes all organismal genes — not just highly expressed ones [13]. Thus, even low-expression genes possess unique codon usage profiles differing from highly expressed genes.

In this study, we propose a novel approach to investigating codon bias ranges through the concept of Genome Subspaces, which define the range of sequence variations across the certain organism. Examining this approach with four distinct test unicellular species have brought captivating observations of separate gene clusters occurring for each species. In addition, we propose that our organism-specific Genome Subspace analysis, owing to its simplicity, will provide new insights into the origin and organization of primitive living organisms and life itself.

## MATERIAL AND METHODS

### Dataset

In this study, we analyzed coding sequence regions from four different species: *Escherichia coli* k12, *Roseiflexus castenholzii* ASM1780v1, *Salinispora tropica* ASM1642v1 and *Saccharomyces cerevisiae* R64-1-1, which were obtained from Ensembl database [https://bacteria.ensembl.org/Escherichia_coli_k_12_gca_004802935/Info/Index, https://bacteria.ensembl.org/Roseiflexus_castenholzii_dsm_13941_gca_000017805/Info/Index, https://bacteria.ensembl.org/Salinispora_tropica_cnb_440_gca_000016425/Info/Index, https://fungi.ensembl.org/Saccharomyces_cerevisiae/Info/Index].

### Gene Space

We analyze gene sequences from the point of token frequencies. Here tokens are short subsequences of a certain fixed length, for example individual nucleotides, codons or codon pairs. To construct the Gene Space, we created n orthogonal axes, each corresponding to the specific token, where *N* is the total number of possible tokens of such length. As we only analyze amino acid-coding codons, *N* = 4 for nucleotides; *N* = 61 for codons; *N* = 3721 for codon pairs. Each axis spans from 0 (in the origin) to 1. For each gene we calculated the number of token occurrences and divided them by the total number of tokens in the gene, thus receiving the token frequencies of the gene. Each gene can be represented as a point, with n coordinates (token frequencies) describing its location on each of the axes. Due to the normalization of the codon frequencies, the gene position range represents an *n* – 1 dimensional simplex with *n* vertices located at the points where a single axis value equals 1. This range was denoted as the Gene Space. Any nucleotide sequence can be represented as a point in the Gene Space with the coordinates described by its token frequencies.

### Separation of genes from two species

In order to separate clusters of genes from two different species we used the Support Vector Machine (SVM) [14] method. It creates an (*n* – 1)-dimensional hyperplane that splits the Gene Space into two parts. When training SVM, we utilized the linear kernel so that the boundary would be a hyperplane.

In order to examine the efficiency of species separation, we split each organism’s gene dataset into two disjoint equal size sets. One of the sets from both organisms was used for model training and the other one was used to calculate the probability of the correct identification of the genes. Then we swapped the sets and repeated the process (trained the model on the second sets and validated on the first ones). The resulting probability value is a mean of the two results.

### Genome Subspace

Genome Subspace is an area of the Gene Space that contains the majority of genes from that organism. In order to define the subspace, we utilized the One Class SVM [15] method as it does not require a negative dataset, which is absent in the unary classification task. The model was trained using a linear kernel and the upper bound for the number of training errors was set to 0.1, so the subspace would contain at least 90% of the genes from the training set. The result is a hyperplane that splits the Gene Space into two parts. One of them contains the majority of genes and is the organism’s Genome Subspace.

### Overfitting ratio

In order to control the level of overfitting for the used models, we randomly split the dataset into two separate sets with the same number of genes in each one. The first set was used for the Genome Subspace model training, and the second was used for testing the prediction accuracy. When training the One Class SVM model, some genes are left outside the subspace, we refer to them as the Training loss. Genes from the testing set that are left outside the subspace are referred to as the Validation loss.

To eliminate any discrepancies due to the specific split of the dataset, the model then was trained on the second set, and tested on the first. The resulting overfitting ratio is calculated as an average of the two measurements.

### Distance between points in the Gene Space

If we know coordinates of the two points in the Gene Space, then the distance between them can be calculated as the Euclidean distance.

The largest possible distance inside the Gene Space is between any two vertices and is equal to 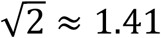. The center point of the Gene Space is located in the geometric center of the simplex and corresponds to the equal distribution of tokens in the nucleotide sequence. The furthest points of the Gene Space from the center point are simplex vertices, and so the maximum distance is 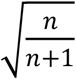, where n is the number of vertices in the Gene Space. Therefore, for codons (*n* = 61) and codon pairs (*n* = 3721) this distance can be considered as 1.

### Subspace volume

Genome Subspace boundary is a hyperplane that divides the Gene Space into two parts. It is located in the relative neighborhood of the center of the Gene Space. The Gene Space is highly dimensional (60 dimensions for codons and 3720 for codon pairs), therefore calculating the volume of the cut off part of the space is highly problematic due to the computational complexity of the process. The most optimal way is to estimate the volume of the cut off part from the number of vertices that fit inside the subspace compared to the total number of simplex vertices. There are more accurate estimates, but due to the unevenness of the gene distribution inside the Gene Space, they don’t make much sense for comparisons between different organisms.

## RESULTS

Any nucleotide sequence can be split into a series of short subsequences — such as individual nucleotides, codons, codon pairs, etc., also called *k*-mers in the literature [16–18]; however, for the sake of simplicity and clarity for the common audience we propose the term token. Consequently, any gene can be characterized by the frequencies of its tokens.

In this study, we introduce the concept of a multidimensional Gene Space capable of describing any possible set of genes. Within this space, a gene is represented as an *n*-dimensional vector, where n denotes the total number of possible tokens (e.g., 4 for nucleotides, 61 for codons, 3721 for codon pairs). For this work, we exclusively use amino acid-encoding codons; thus, any combination containing a stop codon is excluded. Gene Space is based on n orthogonal axes, each corresponding to the frequency of a specific token, with values ranging from 0 to 1, representing the frequency of that token in a given nucleotide sequence normalized to the number of tokens in the sequence. A visual example for the nucleotide-based Gene Space is presented in Fig. 1.

**Figure 1.**
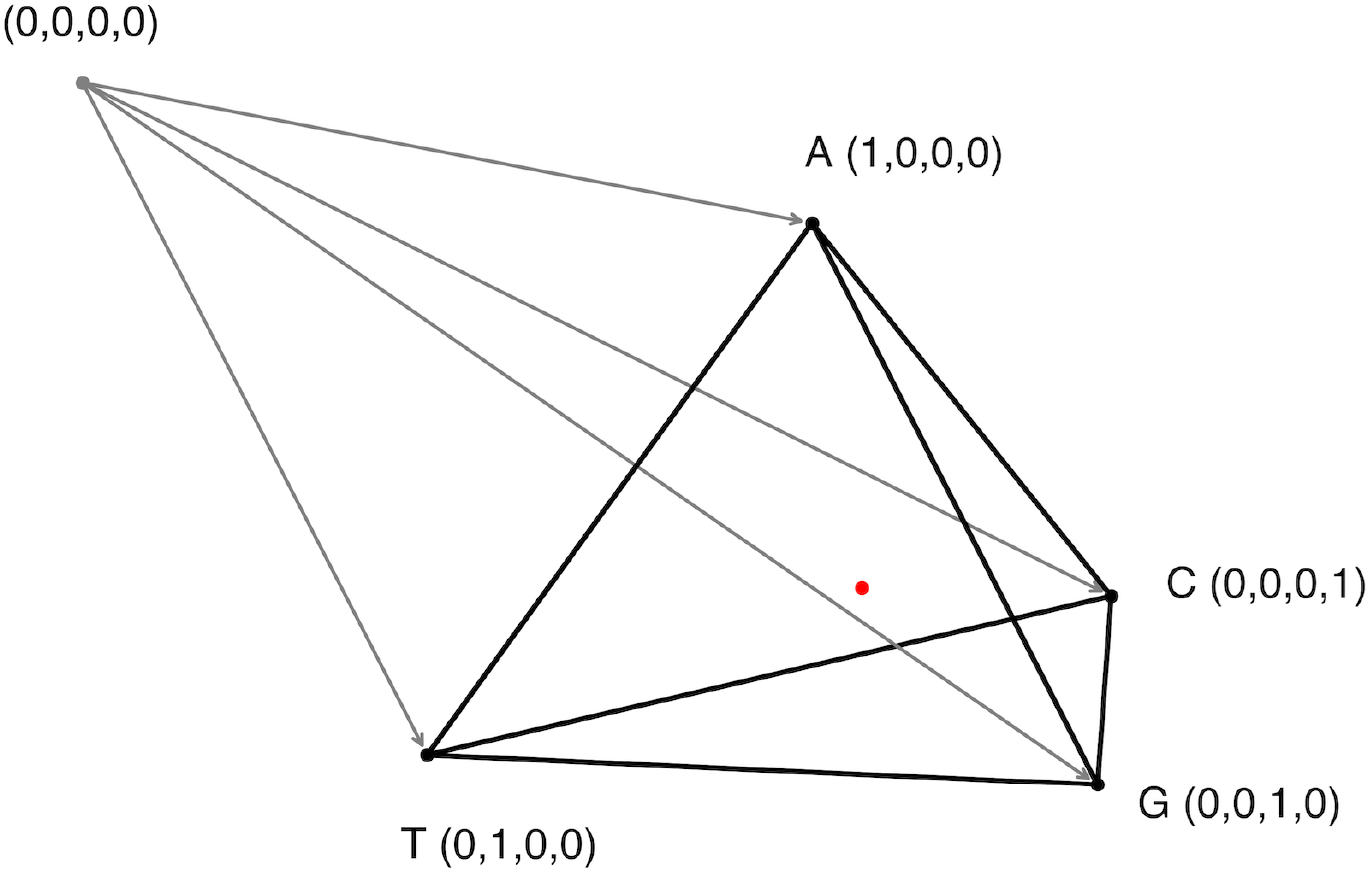
Grey point with coordinates (0, 0, 0, 0) shows the origin and grey arrows represent four orthogonal axes for the four nucleotides (A, T, G, C). Black dots show the points equal to 1 on each of the axes. The wireframe pyramid shows the Gene Space for individual nucleotides. The red dot with coordinates (0.25, 0.25, 0.25, 0.25) shows the geometric center of the pyramid, representing a sequence with equiprobable nucleotides.

Since token frequencies are normalized (summing up to 1), the Gene Space constitutes an (*n* – 1)-dimensional simplex containing n vertices. Each vertex lies at a point where one token’s frequency equals 1 and all others are 0. Thus, Gene Space is a bounded region within *n*-dimensional space, constrained within the interval from 0 to 1 along each axis.

Any gene can be represented in the Gene Space as a point with n coordinates on each of the axes. But due to the token normalization, the gene position range forms an (*n* – 1)-dimensional simplex with *n* vertices at the points, where exactly one of the axes is equal to 1. We designated this limited area as the Gene Space.

Every nucleotide sequence maps to a specific point in Gene Space, yet multiple sequences can occupy identical positions if they share token frequencies. For example: sequences ATGGCA and ATGATGGCAGCA coincide in nucleotide-based and codon-based Gene Spaces (identical codon frequencies) but diverge in codon-pair-based space.

Conversely, ATGGCAATG and ATGGCAATGGCAATG coincide in codon-pair-based space but differ in nucleotide/codon spaces.

To illustrate Gene Space organization, we use four arbitrarily selected unicellular species belonging to evolutionary distant groups (see Table 2) with comparable genome sizes. Among three bacteria, one, *E. coli*, belongs to the kingdom Pseudomonadati, while the two others, *R. castenholzii* and *S. tropica*, shares the same Bacillati kingdom, but belong to different phyla, Chloroflexota and Actinomycetota, respectively. *S. cerevisiae* is eukaryotic fungi. Full phylogeny for the selected organisms is presented in the supplementary file.

Let us consider the simplest nucleotide Gene Space, which can be visualized as a 3-dimensional simplex (tetrahedron) (Fig. 2). Subsequent analyses use higher-dimensional codon and codon-pair Gene Spaces for increased precision, though nucleotide Gene Space remains optimal for intuitive visualization.

**Figure 2.**
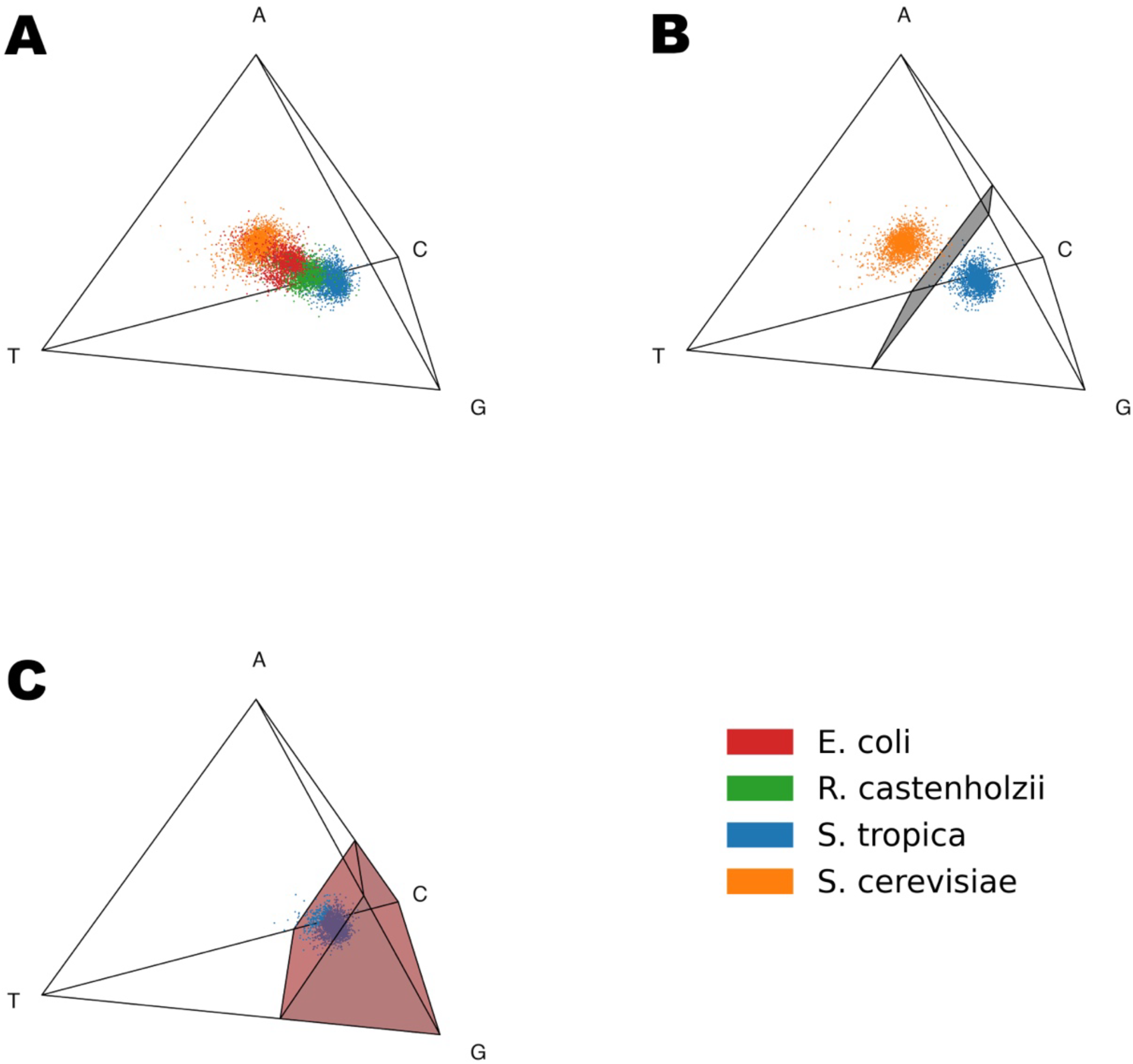
(**A**) A visualization of the 3-dimensional space of the frequencies of nucleotides A, T, G, and C (k = 1). Each point corresponds to a single gene from one of the organisms: orange points represent *Saccharomyces cerevisiae*, red points *Escherichia coli*, green points *Roseiflexus castenholzii*, and blue points *Salinispora tropica*. (**B**) Hyperplane gene separation of different organisms in 4-dimensional space of nucleotide frequencies for *S. cerevisiae* (orange points) and *S. tropica* (blue points). (**C**) *S. tropica* Genome Subspace and separation of its genes by the subspace boundary inside the codon-based Gene Space.

Fig. 2A demonstrates the 3D Gene Space based on the nucleotide frequencies (A, T, G, C). Each point represents a single gene from one of the four studied organisms: *S. cerevisiae* (orange), *E. coli* (red), *R. castenholzii* (green), *S. tropica* (blue). Even inside the simplest nucleotide-based Gene Space, genes form distinct organism-specific clusters.

From Fig. 2A, it is evident that genes from different organisms form distinct clusters and can be separated even at the nucleotide level. In order to do so we used the Support Vector Machine (SVM) method to construct a decision boundary separating a pair of organisms. Fig. 2B shows the decision plane built using the SVM to separate *S. cerevisiae* and *S. tropica* genes in the nucleotide Gene Space. While there is a clear dependency between positions of the genes inside the nucleotide Gene Space and GC content of the organism, there is a lot of variation between the positions of the individual genes, meaning that it is not the only factor influencing the positioning of genes inside the Gene Space. There are also lots of codon combinations that could achieve a similar GC content, which can only be identified by models with longer tokens.

We applied the Support Vector Machine (SVM) method to separate each pair of the test organisms in the nucleotide, codon and codon pair Gene Spaces (Table 1). As shown in Table 1, the increase in Gene Space dimensionality improves the accuracy of gene classification for each pair of organisms. Specifically, in the codon-pair space, the probability of the correct gene to organism matching exceeds 97% for all the organism pairs. Thus, longer tokens that form higher-dimensional Gene Spaces enable more precise compartmentalization of individual organisms. Based on these observations, we hypothesize that each organism occupies a limited region within the Gene Space.

**Table 1.** The probability of correct identification of genes of an organism from a test set of genes when separating two organisms using SVM at three different tokenization levels.

| Organism | Token | *E. coli* | *R. castenholzii* | *S. tropica* | *S. cerevisiae* |
| --- | --- | --- | --- | --- | --- |
| *E. coli* | nucleotides | — | 0.903 | 0.989 | 0.893 |
|  | codons | — | 0.968 | 0.990 | 0.974 |
|  | codon pairs | — | 0.982 | 0.996 | 0.978 |
| *R. castenholzii* | nucleotides | 0.903 | — | 0.910 | 0.979 |
|  | codons | 0.968 | — | 0.981 | 0.985 |
|  | codon pairs | 0.982 | — | 0.989 | 0.991 |
| *S. tropica* | nucleotides | 0.989 | 0.910 | — | 0.998 |
|  | codons | 0.990 | 0.981 | — | 0.998 |
|  | codon pairs | 0.996 | 0.989 | — | 0.998 |
| *S. cerevisiae* | nucleotides | 0.893 | 0.979 | 0.998 | — |
|  | codons | 0.974 | 0.985 | 0.998 | — |
|  | codon pairs | 0.978 | 0.991 | 0.998 | — |

Thus, we have demonstrated that the proposed Gene Space allows for effective and accurate separation of genomes. Based on this, we hypothesize that the genes of each organism occupy a specific bounded region in the Gene Space, which we will refer to as the Genome (organism’s) Subspace. When developing a method for identification of such subspaces within the Gene Space, there are two main constraints: the majority of the organism’s genes should fall within its corresponding subspace and the size of the allocated Genome Subspace should be minimized around the actual genes.

Unlike the binary classification problem of separating genes from two organisms, Genome Subspace identification lacks a negative dataset, thus requiring a different approach. For this task, we employed a one-class Support Vector Machine (one-class SVM), trained to identify a certain percentage (we used 90%) of genes from the reference set. As a result, the one-class SVM produced a boundary of the subspace represented by a hyperplane dividing the Gene Space into two parts, one of which constitutes the organism’s subspace.

Fig. 2C illustrates the Genome Subspace boundary for *S. tropica* in the nucleotide Gene Space: the dark region represents the subspace, dark blue points correspond to genes inside the subspace, and light blue points indicate the 10% of *S. tropica* genes excluded from the subspace.

Thus, we can delineate the boundary and identify the Genome Subspace of a specific organism within the Gene Space. It is important to note that when applying machine learning methods, especially for high-dimensional data, the issue of overfitting inevitably arises. To assess overfitting, we calculated overfitting ratios, defined as the ratio between the model’s loss on disjoint training and validation datasets. An overfitting ratio equal to one indicates that the validation loss matches the training loss, while higher values signify more pronounced overfitting.

The results of this test are presented in Table 2. As observed, for both nucleotide- and codon-based spaces, the overfitting ratios are close to 1, indicating virtually no overfitting. For codon-pair spaces, the values approach 1.3, meaning that compared to the 10% of genes excluded from the Genome Subspaces in the training set, approximately 13% of genes fall outside of the Genome Subspaces in the validation set. However, this level of overfitting remains manageable and should not pose significant issues.

**Table 2.** Overfitting ratios for the Genome Subspace models trained on four different organisms: 90% of genes from each organism are inside the subspace.

| Trained on | Family | Number of genes | Nucleotide | Codon | Codon Pair |
| --- | --- | --- | --- | --- | --- |
| *E. coli* | Prokaryote;<br>Bacteria;<br>Pseudomonadati | 4852 | 1.002 | 1.037 | 1.335 |
| *R. castenholzii* | Prokaryote;<br>Bacteria;<br>Bacillati | 4855 | 1.023 | 1.014 | 1.285 |
| *S. tropica* | Prokaryote;<br>Bacteria;<br>Bacillati | 4666 | 1.006 | 1.024 | 1.141 |
| *S. cerevisiae* | Eukaryota | 6600 | 1.009 | 1.017 | 1.370 |

It is also worth noting that the datasets were split in half for this test, implying that the number of genes in the training sets was smaller than the number of tokens in the codon-pair model. Therefore, training on the full gene set could reduce the model’s level of overfitting, supporting our recommendation to use both codon-based Gene Space and codon-pair-based Gene Space.

The Genome Subspace for each organism represents a specific and unique mode of genome optimization. Identification and comparison of the Genome Subspaces for different organisms can be highly informative. The most straightforward metric for a subspace would be its volume; however, calculating volume in high-dimensional spaces such as those based on codons and codon pairs is computationally challenging due to extreme complexity. Therefore, we propose an estimate for the volume of an organism’s Genome Subspace from the number of the Gene Space vertices *n*_*a*_contained inside of it. These estimates would depend on the total Gene Space size *N*, which equals the number of tokens used to construct it and the variability between genes of a specific organism. The resulting volume estimates for the Genome Subspaces are presented in Table 3.

**Table 3.** Estimated volumes of the Genome Subspaces of the four studied organisms in the Gene Space.

|  | The total size of the Gene Space | *E. coli* | *R. castenholzii* | *S. tropica* | *S. cerevisiae* |
| --- | --- | --- | --- | --- | --- |
| Nucleotides | 4 | 1 | 2 | 2 | 2 |
| Codons | 61 | 20 | 18 | 15 | 20 |
| Codon Pairs | 3721 | 778 | 758 | 375 | 992 |

Table 3 shows that each organism’s Genome Subspace occupies a substantial portion of the entire Gene Space, although genes are unevenly distributed within these subspaces. Since we conceptualize the Gene Space as a simplex — where each vertex corresponds to the sequences entirely composed of a single token (whether it be a nucleotide, codon, or codon pair) — the geometric center of the simplex represents a sequence with all tokens present in equal proportions. Therefore, the position of each gene can be interpreted as the distance from this center point, with the maximum possible distance being the distance to any vertex, which is approximately equal to 1. As the vertices represent sequences composed entirely of a single token, this distance serves as a measure of inequality in token usage within the gene. Fig. 3 illustrates the distributions of distances from each gene of the studied organisms to the center of Gene Space for both codon-based and codon-pair-based Gene Spaces.

**Figure 3.**
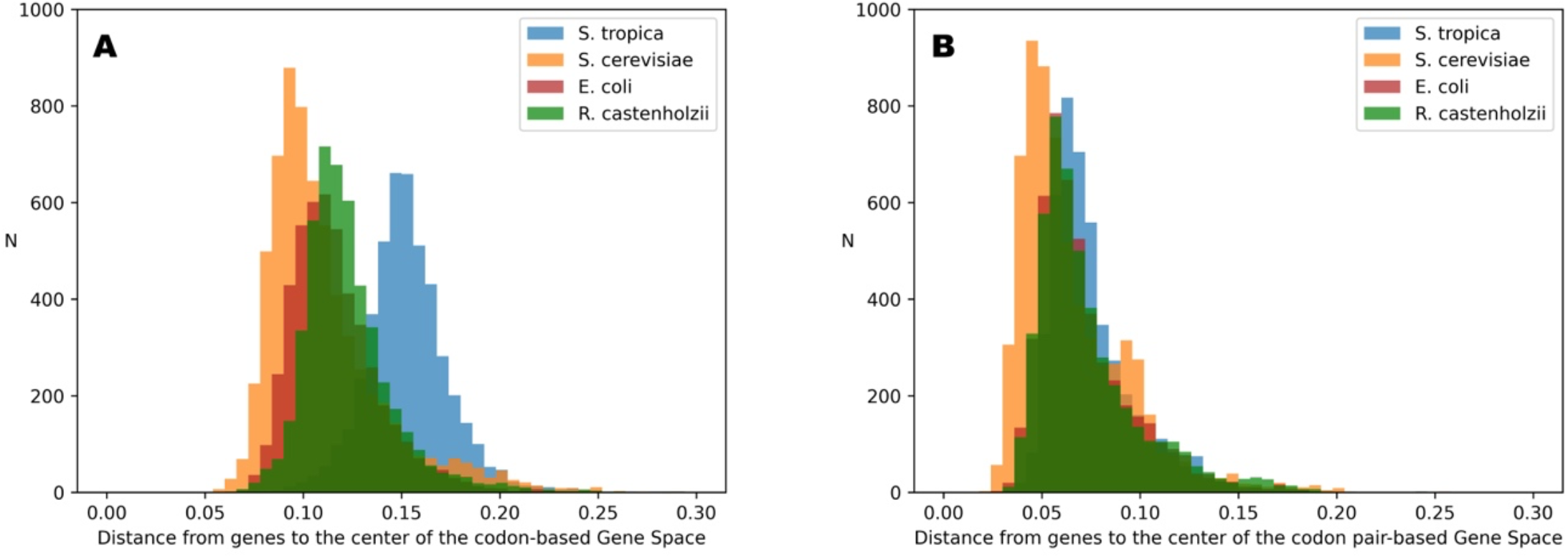
Distributions of distances from genes to the centers of the codon (**A**) and codon pair (**B**) Gene Spaces for the four studied organisms.

Fig. 3 shows that, despite the minor differences, all distributions occupy the same range of distances in the codon-pair space. Moreover, all genes are clustered closer to the center point (0 on the distance scale) than to the vertices, with the most distant gene (i.e., the gene with the most non-random codon-pair composition) located at approximately 20% of the maximum possible distance (equals 1 for the Gene Space vertices). For codon-based spaces, there are greater differences between the distributions of different organisms; however, they still occupy about 20% of the possible distance range (from 5% to 25% from the center point).

This means that although Gene Space is quite sparse and much of it is empty, the gene-occupied region extends at most 25% away from the center point. Nevertheless, a relatively simple machine learning approach can successfully identify token variants specific to particular organisms.

While volume estimates for the Genome Subspace work well for comparisons between different organisms, they should not be interpreted as the proportions of the entire Gene Space.

The differences observed in the distributions shown in Fig. 3 can be explained by variations in genome optimization, differences in amino acid composition, and gene length. These factors influence the minimal possible distance from a gene point to the center of the Gene Space.

Let us examine the commonalities and differences among all the organisms studied. Fig. 4 shows the intersections of the simplex vertices belonging to the Genome Subspace of each of the four studied organisms. It can be seen that 39 vertices in the codon-pair Gene Space are common to all four organisms, while only one vertex in the codon space is shared by all of them (corresponding to glutamic acid). Among the bacteria, seven codons are shared, corresponding to such amino acids as alanine, glycine, isoleucine, leucine, valine, and glutamine.

**Figure 4.**
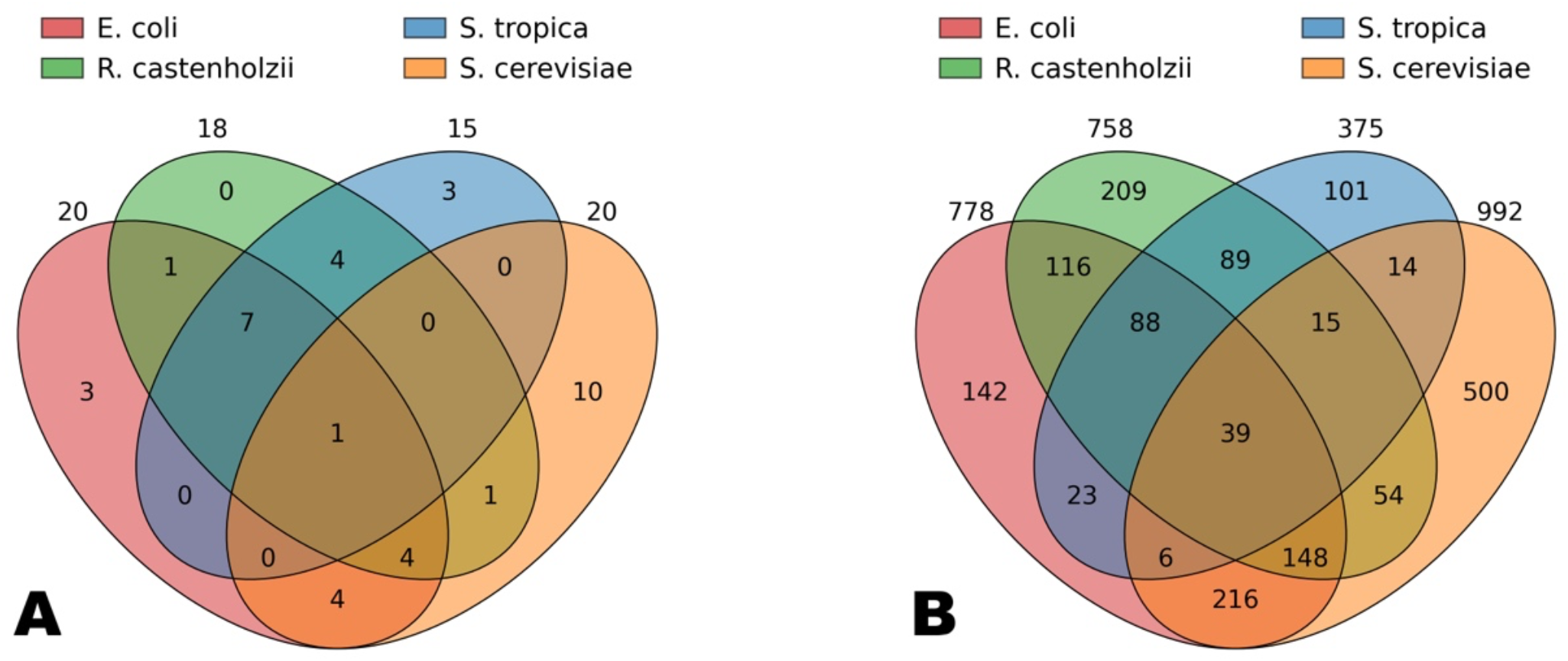
Venn diagrams for the volumes of the Genome Subspace intersections for the four studied organisms in the codon (**A**) and codon pair (**B**) Gene Spaces.

It is important to note that the simplex vertices included in the Genome Subspaces are arranged such that the diversity of amino acids within a Genome Subspace is consistently proportionally higher than the diversity of codons. For example, in the codon Gene Space, each organism’s subspace includes vertices corresponding to more than 10 amino acids, accounting for over half of the total amino acid diversity. In contrast, these codons represent less than one-third of the total codon diversity. This suggests that Genome Subspaces tend to encompass the maximal or necessary number of amino acids while preserving the unique codon composition of each organism.

Notably, cysteine and tryptophan vertices are not included in any of the studied subspaces, and histidine, serine, and tyrosine are absent from the bacterial Genome Subspaces. Meanwhile, amino acids such as alanine, aspartic acid, glutamic acid, isoleucine, and leucine are present in the Genome Subspaces of all four organisms, often each encoded by their organism-specific codons.

Furthermore, the Venn diagrams presented in Fig. 4 allow conclusions to be drawn about the evolutionary relationships among the studied organisms.

The distribution of distances from each gene to the center of Gene Space (see Fig. 3) reveals non-uniform density and suggests genomic organization into primary and secondary genes within this spatial context. To quantify the subspace filling uniformity, we artificially constrained Genome Subspace volume so that it would fit only a fraction of the organism’s genes, and estimated the subspace volumes. Results (Fig. 5) demonstrate a nonlinear dependence, confirming the unevenness of the gene distributions. Vertex analysis across *v* values showed an invariant composition in >95% of cases, validating Genome Subspaces as a robust, organism-specific genomic characteristic.

**Figure 5.**
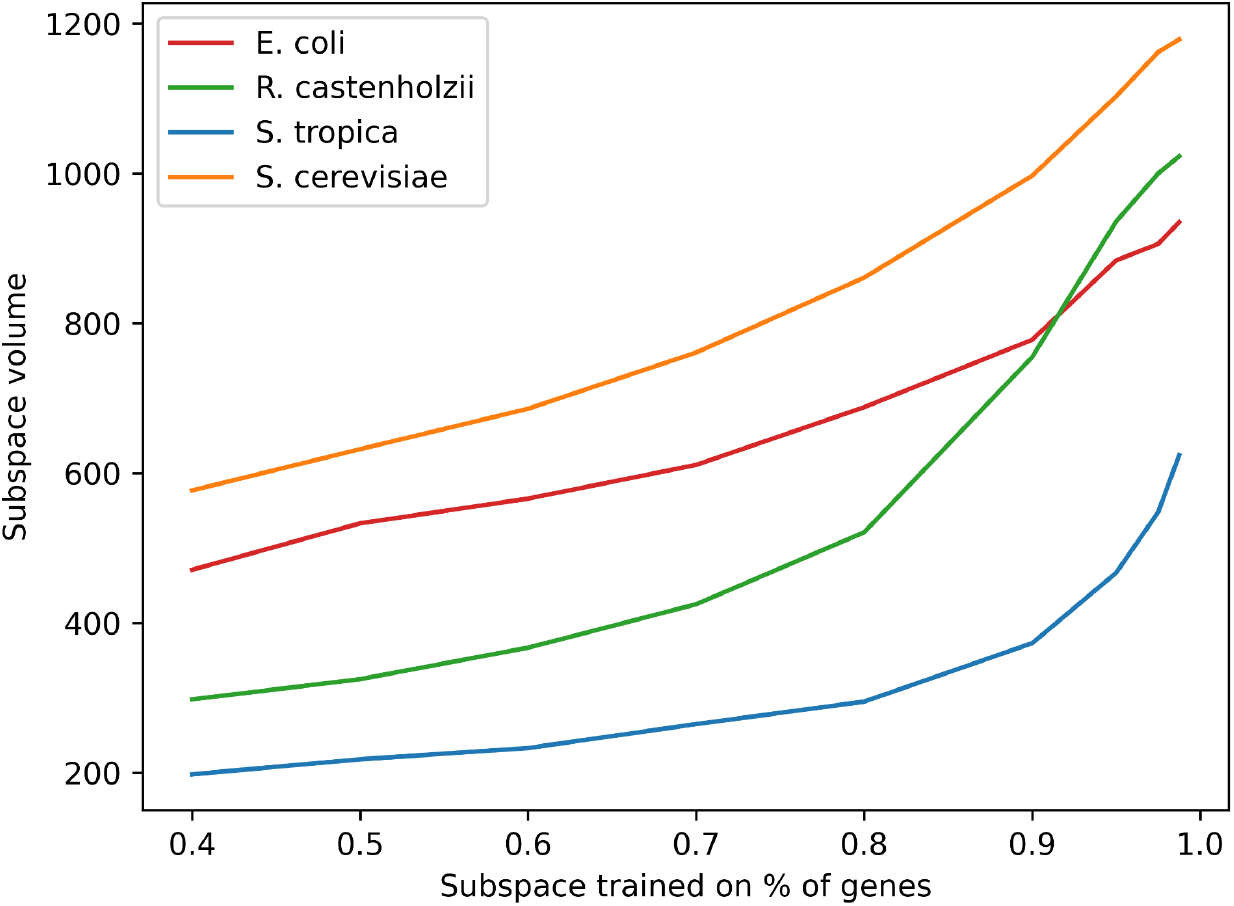
Dependence of the volume of Genome Subspaces of organisms on the percentage of genes of this organism falling into the distribution core.

## DISCUSSION

The fundamental law of life dictates that all genes of an organism are encoded using the same universal principle: each amino acid can be specified by 1 to 6 synonymous codons. As a result, each amino acid sequence can be represented by an astronomically large number of synonymous nucleotide sequences. But despite all the diversity, organisms select certain configurations that are unique for each species due to the different optimization paths [19,20]. This phenomenon is well known and researched and the most popular index that describes it is the codon bias [1,21], which shows codon preferences of the organism.

In this study, we introduced the concept of a multidimensional Gene Space based on the normalized frequencies of the nucleotide tokens — ranging from single nucleotides to codons and codon pairs — and demonstrated its utility for characterizing and distinguishing genes across different organisms. Our results show that genes from each species occupy a distinct, bounded region within the Gene Space, which we define as an organism-specific confined Genome Subspace. This finding supports the hypothesis that each organism’s genome is optimized as a whole in a unique manner, being reflected in the specific distribution of token frequencies that shape its subspace. Codon-based Genome Subspaces practically reflect the codon bias range, showing not only the codon bias preferred by the organism, but also the possible variations in codon usage across the genes.

The application of machine learning methods, particularly the One-class Support Vector Machine (OcSVM) [15], allowed us to effectively delineate these subspaces especially in the high-dimensional codon and codon-pair Gene Spaces. The classification accuracy improved with the increase in dimensionality, but the method becomes extremely computationally complex for tokens longer than six nucleotides due to the computational complexity. Importantly, overfitting tests revealed that our models generalize well, with overfitting ratios close to one for nucleotide and codon Gene Spaces, and still manageable levels for the codon-pair Gene Spaces. This robustness justifies the use of both codon and codon-pair Gene Spaces for genome analysis.

While longer tokens allow for more effective genome separation, there is a practical limit to the length. With longer tokens there are more possible token variations and so, the Gene Space contains even more dimensions. For example, for 9 nucleotide long tokens the Gene Space would be 226980-dimensional, significantly exceeding the number of genes in any living organism. And if a Genome Subspace model was to be trained in such Gene Space, it would suffer from extreme overfitting. Thus, we limited ourselves to codon and codon pair-based models.

Volume determination for these subspaces, while challenging due to the computational complexity of high-dimensional spaces, was approached as an estimate from the number of simplex vertices included in each organism’s Genome Subspace. This proxy measure revealed that subspaces cover substantial portions of the overall Gene Space but are unevenly filled by genes. Analysis of distances from the center of the Gene Space — a point representing equal token usage — indicated that all genes cluster closer to the center than to vertices, suggesting that genomes maintain a balance between variation and specificity in token usage. The maximum observed distance was about 20–25% of the total possible distance, reflecting constraints imposed by the amino acid composition and gene length. The furthest points from the Gene Space center are the vertices that represent sequences consisting of a single token. Therefore, the distance from a gene to the Gene Space center shows the degree of heterogeneity in the token distribution, linked to the optimization of the organism’s genes for its life conditions [22,23].

Comparative analysis of the Genome Subspace intersections highlighted both shared and unique features among the organisms. For example, while 39 vertices were common to all four studied species in the codon-pair Gene Space, only one vertex was shared by all four studied organisms in codon Gene Space. Bacterial subspaces shared codons corresponding to a core set of amino acids, indicating evolutionary conservation, whereas other amino acids were organism-specific. This pattern suggests that subspaces strive to encompass a broad and necessary set of amino acids while preserving distinct codon usage profiles, reflecting both functional requirements and evolutionary history.

Finally, investigation of gene distribution density within the Genome Subspaces revealed a nonlinear relationship between the fraction of genes included and the occupied volume, confirming that gene distribution is heterogeneous rather than uniform. This heterogeneity likely reflects some biological factors such as gene expression levels, functional clustering, and evolutionary pressures.

Overall, the Gene Space framework and associated machine learning approaches provide a novel and powerful approach for understanding genome organization, optimization, and evolution. By capturing the complex interplay of token usage patterns, gene length, and amino acid composition, this approach offers new opportunities for comparative genomics and synthetic biology, including improved gene optimization strategies tailored to the specific organisms.

## ACKNOWLEDGEMENTS

We are grateful to Maria Yurkova and Dmitrii Kostenko for their seminar discussion and support.

## AUTHOR CONTRIBUTIONS

Konstantin Zaytsev: Conceptualization, Methodology, Software, Validation, Formal analysis, Investigation, Data curation, Visualization, Writing — original draft preparation, Writing — review & editing. Natalia Bogatyreva: Methodology, Validation, Formal analysis, Data curation, Visualization, Writing — original draft preparation, Writing — review & editing. Alexey Fedorov: Conceptualization, Validation, Resources, Writing — review & editing, Supervision, Project administration, Funding acquisition.

## DATA AVAILABILITY

Python module for the Gene Space analysis is available at https://github.com/conzaytsev/GeneSpace

## REFERENCES

1. Grantham, R.; Gautier, C.; Gouy, M.; Mercier, R.; Pavé, A. Codon Catalog Usage and the Genome Hypothesis. Nucleic Acids Res. 1980, 8, 197–197, doi:10.1093/nar/8.1.197-c.

2. Plotkin, J.B.; Kudla, G. Synonymous but Not the Same: The Causes and Consequences of Codon Bias. Nat. Rev. Genet. 2011, 12, 32–42, doi:10.1038/nrg2899.

3. Lynn, D.J. Synonymous Codon Usage Is Subject to Selection in Thermophilic Bacteria. Nucleic Acids Res. 2002, 30, 4272–4277, doi:10.1093/nar/gkf546.

4. Chen, S.L.; Lee, W.; Hottes, A.K.; Shapiro, L.; McAdams, H.H. Codon Usage between Genomes Is Constrained by Genome-Wide Mutational Processes. Proc. Natl. Acad. Sci. 2004, 101, 3480–3485, doi:10.1073/pnas.0307827100.

5. Muto, A.; Osawa, S. The Guanine and Cytosine Content of Genomic DNA and Bacterial Evolution. Proc. Natl. Acad. Sci. 1987, 84, 166–169, doi:10.1073/pnas.84.1.166.

6. Ikemura, T. Correlation between the Abundance of Escherichia Coli Transfer RNAs and the Occurrence of the Respective Codons in Its Protein Genes. J. Mol. Biol. 1981, 146, 1–21, doi:10.1016/0022-2836(81)90363-6.

7. Ikemura, T. Correlation between the Abundance of Escherichia Coli Transfer RNAs and the Occurrence of the Respective Codons in Its Protein Genes: A Proposal for a Synonymous Codon Choice That Is Optimal for the E. Coli Translational System. J. Mol. Biol. 1981, 151, 389–409, doi:10.1016/0022-2836(81)90003-6.

8. Frumkin, I.; Lajoie, M.J.; Gregg, C.J.; Hornung, G.; Church, G.M.; Pilpel, Y. Codon Usage of Highly Expressed Genes Affects Proteome-Wide Translation Efficiency. Proc. Natl. Acad. Sci. 2018, 115, doi:10.1073/pnas.1719375115.

9. Hoffmann, F.; Rinas, U. Stress Induced by Recombinant Protein Production in Escherichia Coli. In Physiological Stress Responses in Bioprocesses; Advances in Biochemical Engineering/Biotechnology; Springer Berlin Heidelberg: Berlin, Heidelberg, 2004; Vol. 89, pp. 73–92 ISBN 978-3-540-20311-7.

10. Huang, Y.; Lin, T.; Lu, L.; Cai, F.; Lin, J.; Jiang, Y.; Lin, Y. Codon Pair Optimization (CPO): A Software Tool for Synthetic Gene Design Based on Codon Pair Bias to Improve the Expression of Recombinant Proteins in Pichia Pastoris. Microb. Cell Factories 2021, 20, 209, doi:10.1186/s12934-021-01696-y.

11. Powell, J.R.; Moriyama, E.N. Evolution of Codon Usage Bias in Drosophila. Proc. Natl. Acad. Sci. 1997, 94, 7784–7790, doi:10.1073/pnas.94.15.7784.

12. Liu, Y.; Yang, Q.; Zhao, F. Synonymous but Not Silent: The Codon Usage Code for Gene Expression and Protein Folding. Annu. Rev. Biochem. 2021, 90, 375–401, doi:10.1146/annurev-biochem-071320-112701.

13. Zaytsev, K.; Bogatyreva, N.; Fedorov, A. Link Between Individual Codon Frequencies and Protein Expression: Going Beyond Codon Adaptation Index. Int. J. Mol. Sci. 2024, 25, 11622, doi:10.3390/ijms252111622.

14. Boser, B.E.; Guyon, I.M.; Vapnik, V.N. A Training Algorithm for Optimal Margin Classifiers. In Proceedings of the Proceedings of the fifth annual workshop on Computational learning theory; ACM: Pittsburgh Pennsylvania USA, July 1992; pp. 144–152.

15. Schölkopf, B.; Williamson, R.C.; Smola, A.; Shawe-Taylor, J.; Platt, J. Support Vector Method for Novelty Detection. In Proceedings of the Advances in Neural Information Processing Systems; Solla, S., Leen, T., Müller, K., Eds.; MIT Press, 1999; Vol. 12.

16. Moeckel, C.; Mareboina, M.; Konnaris, M.A.; Chan, C.S.Y.; Mouratidis, I.; Montgomery, A.; Chantzi, N.; Pavlopoulos, G.A.; Georgakopoulos-Soares, I. A Survey of K-Mer Methods and Applications in Bioinformatics. Comput. Struct. Biotechnol. J. 2024, 23, 2289–2303, doi:10.1016/j.csbj.2024.05.025.

17. Smith, R.P.; Riesenfeld, S.J.; Holloway, A.K.; Li, Q.; Murphy, K.K.; Feliciano, N.M.; Orecchia, L.; Oksenberg, N.; Pollard, K.S.; Ahituv, N. A Compact, in Vivo Screen of All 6-Mers Reveals Drivers of Tissue-Specific Expression and Guides Synthetic Regulatory Element Design. Genome Biol. 2013, 14, doi:10.1186/gb-2013-14-7-r72.

18. Ghandi, M.; Lee, D.; Mohammad-Noori, M.; Beer, M.A. Enhanced Regulatory Sequence Prediction Using Gapped K-Mer Features. PLoS Comput. Biol. 2014, 10, e1003711, doi:10.1371/journal.pcbi.1003711.

19. Tuller, T.; Waldman, Y.Y.; Kupiec, M.; Ruppin, E. Translation Efficiency Is Determined by Both Codon Bias and Folding Energy. Proc. Natl. Acad. Sci. 2010, 107, 3645–3650, doi:10.1073/pnas.0909910107.

20. Quax, T.E.F.; Claassens, N.J.; Söll, D.; van der Oost, J. Codon Bias as a Means to Fine-Tune Gene Expression. Mol. Cell 2015, 59, 149–161, doi:10.1016/j.molcel.2015.05.035.

21. Parvathy, S.T.; Udayasuriyan, V.; Bhadana, V. Codon Usage Bias. Mol. Biol. Rep. 2022, 49, 539–565, doi:10.1007/s11033-021-06749-4.

22. Korotkov, E.; Zaytsev, K.; Fedorov, A. Use of 6 Nucleotide Length Words to Study the Complexity of Gene Sequences from Different Organisms. Entropy 2022, 24, 632, doi:10.3390/e24050632.

23. Botzman, M.; Margalit, H. Variation in Global Codon Usage Bias among Prokaryotic Organisms Is Associated with Their Lifestyles. Genome Biol. 2011, 12, R109, doi:10.1186/gb-2011-12-10-r109.

