## Supplementary figures and images for "Solving the gene classification problem with a novel Gene Space approach"

### supplementary file

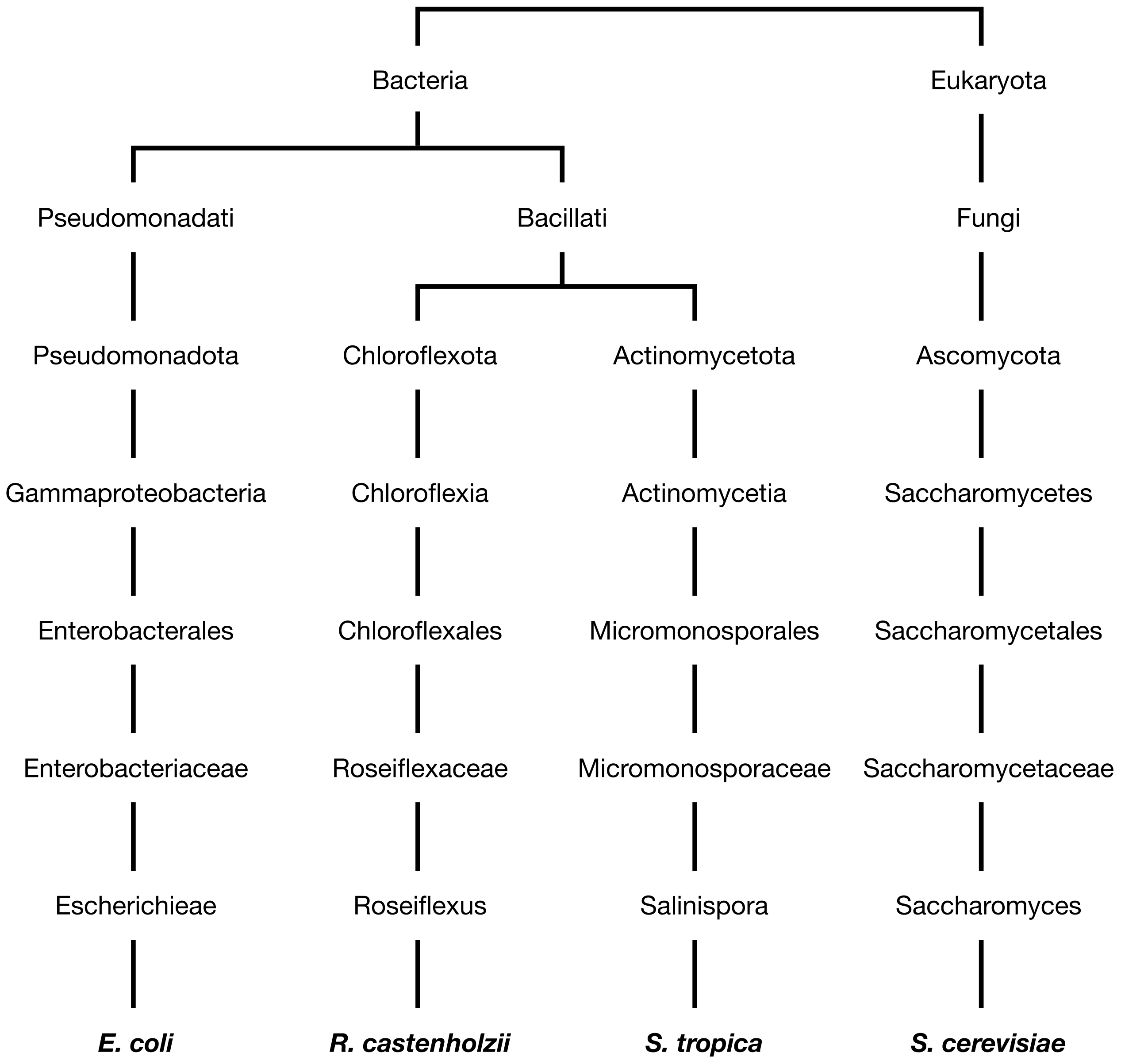
